# Phencyclidine-induced psychosis causes hypersynchronization and disruption of connectivity within prefrontal-hippocampal circuits that is rescued by antipsychotic drugs

**DOI:** 10.1101/2021.02.03.429582

**Authors:** Cristina Delgado-Sallent, Pau Nebot, Thomas Gener, Melina Timplalexi, Amanda B Fath, M Victoria Puig

**Author notes:** **Correspondence:** M. Victoria Puig.

## Abstract

Neural synchrony and functional connectivity are disrupted in neuropsychiatric disorders such as schizophrenia. However, these alterations and how they are affected by commonly prescribed neuropsychiatric medication have not been characterized in depth. Here, we investigated changes in neural dynamics of circuits involving the prefrontal cortex and the hippocampus during psychosis induced by the NMDAR antagonist phencyclidine and subsequent recovery by three different antipsychotic drugs (APDs), the classical APD haloperidol and two atypical APDs, clozapine and risperidone, in freely moving mice. We found that the psychotomimetic effects of phencyclidine were associated with hypersynchronization and disrupted communication of prefrontal-hippocampal pathways. Major alterations occurred in the prefrontal cortex, where phencyclidine increased oscillatory power at delta, high gamma and high frequencies (<100 Hz) and generated aberrant cross-frequency coupling, suggesting the presence of hypersynchronous cortical microcircuits. Cross-regional coupling and phase coherence were also enhanced, further reflecting that the circuit’s functional connectivity was increased. Phencyclidine also redirected the intrinsic flow of information at theta frequencies that traveled from the hippocampus to the prefrontal cortex into delta rhythms that traveled in the opposite direction. The three APDs rescued most phencyclidine-induced changes in power, coupling, phase coherence, and directionality, suggesting common cellular mechanisms of antipsychotic action. However, some differential effects were identified, likely resulting from the distinct affinity the three APDs have for dopamine and serotonin receptors. We therefore investigated how serotonin 1A (5-HT_1A_R) and 2A receptors (5-HT_2A_R) compare to the actions of the APDs. 5-HT_2A_R antagonism by M100907 and 5-HT_1A_R agonism by 8-OH-DPAT rescued phencyclidine-induced increased power, coupling and phase coherence but were unable to normalize the circuit’s theta directionality. This suggests that other targets of the AAPDs working in tandem with 5-HT_1A_Rs and 5-HT_2A_Rs are required to ameliorate this key feature of the circuit.

## INTRODUCTION

Disruption of neural synchrony and spatio-temporal communication in brain circuits involving the prefrontal cortex (PFC) and the hippocampus (HPC) has been suggested to be a hallmark characteristic of neuropsychiatric disorders such as schizophrenia (Baker et al., 2014; Godsil et al., 2013; Sigurdsson and Duvarci, 2016). Schizophrenia patients show abnormal gamma oscillatory activities (>25 Hz) in cortical regions during resting states and sensory processing that have been associated with positive symptoms, such as hallucinations (Grützner et al., 2013; Spencer et al., 2009, 2008; Uhlhaas and Singer, 2010). Moreover, they display disrupted functional connectivity within prefrontal-hippocampal circuits that correlate with poor executive functioning such as working memory and inhibitory control (Benetti et al., 2009; Heckers and Konradi, 2010; Minzenberg et al., 2009; Sigurdsson and Duvarci, 2016).

Rodent models of schizophrenia also show prefrontal-hippocampal dysfunction including increased power of gamma and high frequency oscillations (HFO) (Hunt and Kasicki, 2013; Jodo, 2013) and reduced circuit synchrony in the theta and gamma frequencies (Sigurdsson, 2016; Sigurdsson et al., 2010). Hypofunction of the N-Methyl-D-aspartate receptor (NMDAR) has been proposed to be an important factor in the pathophysiological features of schizophrenia. NMDAR antagonists are used to model schizophrenia in humans and animals, mimicking some of the core positive, negative and cognitive symptoms of the disorder. For example, acute administration of the NMDAR antagonist ketamine produces transient psychotic symptoms in healthy people while it exacerbates pre-existing symptoms in schizophrenia patients (Krystal et al., 1994; Lahti et al., 1995). In rodents, acute NMDAR antagonism mediated by phencyclidine (PCP) or ketamine produces atypical gamma oscillatory activities similar to schizophrenia patients and it also increases the power of HFO (130-180 Hz) across different brain regions that include the PFC and the HPC (Hunt et al., 2015, 2006). However, the exact involvement of prefrontal-hippocampal circuitry in the psychosis induced by NMDAR antagonists has not been described.

The cellular mechanisms of antipsychotic medication used to treat or prevent psychosis are also not fully understood. In fact, how antipsychotic drugs (APDs) affect prefrontal-hippocampal pathways during psychosis is not known. APDs are classified in classical and atypical based on their affinities for dopamine and serotonin receptors. Atypical antipsychotic drugs (AAPDs) primarily bind to serotonin receptors and have been shown to ameliorate cognitive and negative symptoms in addition to psychosis in schizophrenia patients and NMDAR hypofunction models of schizophrenia (Grayson et al., 2007; Houthoofd et al., 2008; Meltzer and McGurk, 1999; Neill et al., 2010). By contrast, classical APDs like haloperidol primarily block dopamine D2 receptors (D2R) and weakly bind to serotonin receptors, which in turn has shown to be less effective in the recovery of cognitive and negative symptoms (Meltzer and Massey, 2011; Meltzer and Huang, 2008). Serotonin 1A (5-HT_1A_R) and serotonin 2A (5-HT_2A_R) receptors are major targets for AAPDs such as risperidone or clozapine. Specifically, 5-HT_2A_R antagonism and 5-HT_1A_R agonism may be linked to AAPDs ability to treat cognitive and negative symptoms (Meltzer and Huang, 2008; Roth et al., 2004). In addition to its serotonergic targets, risperidone also acts as a D2R antagonist, therefore an interaction between the serotonin and dopamine systems may be essential for some AAPD therapeutic actions (Wang et al., 2018). The PFC and HPC densely express serotonin 5-HT_1A_Rs and 5-HT_2A_Rs (Berumen et al., 2012; Celada et al., 2013; Puig and Gener, 2015; Puig and Gulledge, 2011), whereas D2R are moderately present (Puighermanal et al., 2015; Santana et al., 2009). Therefore, APDs binding to these receptors are expected to have major effects on prefrontal-hippocampal neural dynamics.

We have recently shown that risperidone alone reduces local and cross-regional synchronization of prefrontal-hippocampal circuits in freely moving mice, with 5-HT_1A_R, 5-HT_2A_R and D2R shaping distinct frequency bands (Gener et al., 2019). Here we investigate the neural dynamics of prefrontal-hippocampal circuits under the psychotomimetic conditions produced by an acute dose of PCP. We further explore the ability of the APDs risperidone, clozapine and haloperidol, the 5-HT_2A_R antagonist M100907 and the 5-HT_1A_R agonist 8-OH-DPAT to rescue the circuit dysfunction produced by PCP.

## RESULTS

### Phencyclidine boosts prefrontal-hippocampal hypersynchronization and disrupts circuit communication

To elicit psychotomimetic conditions in freely moving mice we administered PCP acutely (10 mg/kg, S.C.; n = 13 mice), a non-competitive antagonist of NMDARs widely used to model psychosis. We performed additional control experiments with saline (n = 10 mice). We investigated how acute PCP influences behavior and neural activity in the prelimbic PFC and the CA1 region of the HPC. We examined the time course of PCP’s effects at 15, 30 and 60 minutes after the injection (Fig. 1a). As previously reported (Lee et al., 2017), PCP induced psychotomimetic symptoms that included catalepsy, stereotypies, head twitches and stiffness of the tail. It also caused ataxia, impaired motor function (i.e., coordination and stability while walking) and reduced general activity for at least 1 hour after its administration (variance of the accelerometer values, Acc: F_1,12_ = 42.77, p <0.0005, repeated measures ANOVA for time [baseline, 15, 30, 60 min after PCP]).

**Figure 1.**
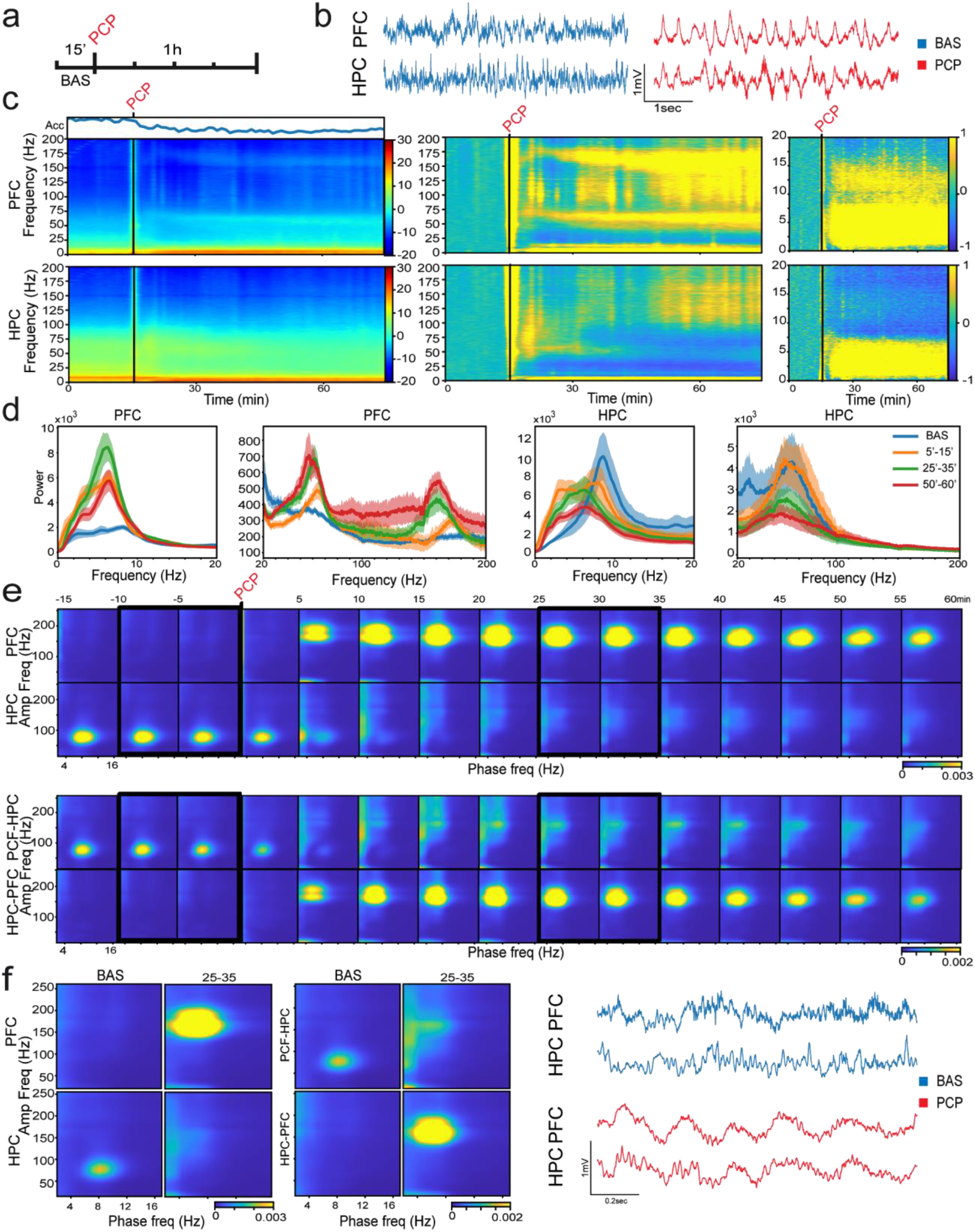
Acute administration of PCP boosts the synchronization of prefrontal and hippocampal microcircuits. **(a)** Experimental protocol. Recording LFP signals in freely moving mice during baseline conditions (BAS) for 15 minutes and for 1 hour following the administration of PCP (SC, 10 mg/kg; n = 13 mice). **(b)** Representative examples of 5 second LFP traces recorded in the PFC and the HPC during baseline (blue) and after the administration of PCP (red) in one mouse. **(c)** Averaged spectrograms of signals in the PFC (upper panels) and the HPC (lower panels). Left panels represent the raw data with the corresponding quantification of the animals’ mobility (Acc, variance of signals from the accelerometer); middle and right panels represent the z-scores relative to baseline. Note the different frequencies on the y-axis between spectrograms. **(d)** Power spectra of PFC (left panels) and HPC (right panels) signals during baseline (blue), 5 to 15 min (orange), 25 to 35 min (green) and 50 to 60 minutes (red) after the administration of PCP. The plots have been divided into 0-20 Hz and 20-200 Hz to facilitate the comparison. **(e)** Comodulation maps quantifying cross-frequency coupling in consecutive non-overlapping 5-min epochs. The x-axis represents phase frequencies (2-16 Hz) and the y-axis represent amplitude frequencies (10-250 Hz). Numbers on top of the panels indicate the minute after PCP administration. (Top) Local modulation index in the PFC and the HPC. (Bottom) Cross-regional modulation index between the PFC phase and the HPC amplitude (upper panels) and between the HPC phase and the PFC amplitude (bottom panels). PCP administration is marked with an arrow. **(f)** Averaged comodulation maps of local and cross-regional modulation index. (Left) Modulation index averaged over 10 minutes during baseline (−10-0 minutes during BAS) and during the 25-35 minutes after PCP administration (marked in panel **e** with black squares). (Right) Representative examples of 1 second LFP signals during baseline (blue) and 30 minutes after the administration of PCP (red). Note the increase of fast delta oscillations (5-6 Hz) and the enhanced local and cross-regional delta-HFO coupling following PCP administration.

This atypical behavior was accompanied by substantial alterations in neural oscillations, particularly in the PFC, that could be readily seen in the unprocessed local field potentials (Fig. 1b). Several aberrant bands emerged in the PFC after injection of PCP: a wide band from 1 to 8 Hz that included the standard “slow” delta (2-5 Hz) and an aberrant “fast” delta (4-8 Hz, peak at 6.4 ± 0.2 Hz), an alpha band (10-14 Hz, peak at 10.2 ± 0.06), a narrow band within the high gamma range (40-75 Hz, peak at 60.3 ± 1.1 Hz), and broadly increased HFOs (100-200 Hz) that included a prominent band between 140-180 Hz (peak at 160.5 ± 2.54 Hz; F_1,12_ = 15.74, 5.42, 23.15, 8.2 and 19.85, respectively; p <0.0005, 0.028, and <0.0005, one-way ANOVA; significant differences with saline controls, two-way ANOVA; Fig. 1c,d and Supplementary Fig. S1). These bands appeared *de novo* and revealed strong hypersynchronization of prefrontal microcircuits by PCP. This enhanced synchronization unstabilized beta waves (18-25 Hz), which was reflected as a reduction in their power (F_1,12_ = 14.67, p <0.0005; Fig. 1c,d). In the HPC, the power changes were modest and mostly differed from the PFC, except for the delta and beta bands that closely followed the patterns found in the PFC (F_1,11_ = 12.86 and 5.15; p <0.0005 and 0.041, respectively). PCP decreased the power of hippocampal theta oscillations (8-12 Hz) with respect to baseline (F_1,11_ = 7.21, p = 0.011), most likely reflecting the reduced mobility of mice. Other differences between power changes in the HPC in comparison with the PFC included the lack of change in the alpha band, a high gamma increase that appeared earlier and was shorter than in the PFC (the first 15 min only; F_1,11_ = 8.48, p = 0.012) and HFOs that emerged much later than in the PFC (40 min after PCP injection; p = 0.04; Fig. 1c,d).

To further assess local and circuit synchronization we quantified cross-frequency phase-amplitude coupling (PAC) between slow rhythms (2-16 Hz) and faster oscillations (50-250 Hz) (Bruns & Eckhorn 2004; Jensen and Colgin, 2007; Scheffer-Teixeira and Tort, 2018). Cross-frequency coupling was investigated both within the PFC and the HPC (local PAC: PFC_phase_-PFC_amp_ and HPC_phase_-HPC_amp_) and at the circuit level via coupling of HPC phase with PFC amplitude and vice versa (circuit PAC: HPC_phase_-PFC_amp_ and PFC_phase_-HPC_amp_). As reported previously (Colgin, 2015; Hentschke et al., 2007; Scheffer-Teixeira and Tort, 2017), we detected the presence of theta-gamma PAC within the CA1 region during baseline periods, with phases varying from 5 to 12 Hz and amplitude frequencies varying from 50 to 100 Hz (Fig. 1e,f). After PCP, this theta-gamma coordination rapidly subsided and delta-HFO coupling emerged (including slow and fast delta modes; [decrease theta-gamma, increase delta-HFO PAC] F_1,10_ = 16.14, 7.82, p = 0.001, 0.005, repeated measures ANOVA; Fig. 1f). In addition, PCP generated PAC within the PFC between fast delta-high gamma and fast delta-HFO (F_1,11_ = 5, 10.69, p = 0.006, <0.0005, respectively; Figure 1e,f). At a circuit level, we found that the phase of PFC theta modulated the amplitude of CA1 high gamma (PFC_phase_-HPC_amp_), as reported recently (Tavares and Tort, 2020; Zhang et al., 2016)(Fig. 1e,f). Again, PCP weakened this circuit coupling while upregulating delta-HFO PAC ([decrease theta-gamma, increase delta-HFO PAC] F_1,10_ = 18.51, 10.235, p = 0.001, 0.001; Figure. 1e,f), as observed in the HPC. Furthermore, PCP promoted fast delta-HFO HPC_phase_-PFC_amp_ comodulation (F_1,10_ = 6.54, p = 0.002; Figure 1e,f) with a pattern closely resembling aberrant prefrontal PAC. We conclude that PCP-induced *de novo* oscillatory activities dominated local and circuit cross-frequency coupling disrupting the theta-gamma coupling inherent to the circuit.

We next investigated PCP-induced alterations in the functional connectivity of the circuit. We first examined prefrontal-hippocampal phase coherence via the weighted phase-lag index (wPLI), a measure that estimates phase synchronization between areas reducing the effects of volume conduction and confounding signals coming from common inputs (Hardmeier et al., 2014; Vinck et al., 2011). We computed wPLI in consecutive one-second time windows to build PLI spectrograms that allowed us to better assess changes of phase coherence over time. These analyses revealed a circuit hypersynchronization at fast frequencies after administration of PCP (high gamma and HFO wPLI: F_1,11_ = 2.65, 9.95, p = 0.065, <0.0005, respectively), while phase coherence at middle frequencies decreased (theta, alpha and beta wPLI: F_1,11_ = 14.02, 12.27, 6.94, p <0.0005 and 0.015, respectively; significant differences with saline controls). Moreover, the normal synchronization at theta shifted to fast delta (peak from 8.45 ± 0.15 to 6.36 ± 0.43) for at least 45 min after PCP injection (Fig. 2a). We also quantified the directionality of the flow of information between the PFC and the HPC via the phase slope index or PSI (HPC-to-PFC or PFC-to-HPC directionality) (Nolte et al., 2008). We computed the PSI in consecutive one-second time windows to build PSI spectrograms that allowed us to keep track of directionality changes over time after PCP induction. We found that during baseline periods, a strong flow of information at theta originated in the HPC and traveled to the PFC. This theta connectivity was also disrupted by PCP (F_1,11_ = 4.36, p = 0.011) by shifting the frequency to fast delta and reversing its directionality from the PFC to the HPC (F_1,11_ = 6.03, p = 0.013; Fig. 2b).

**Figure 2.**
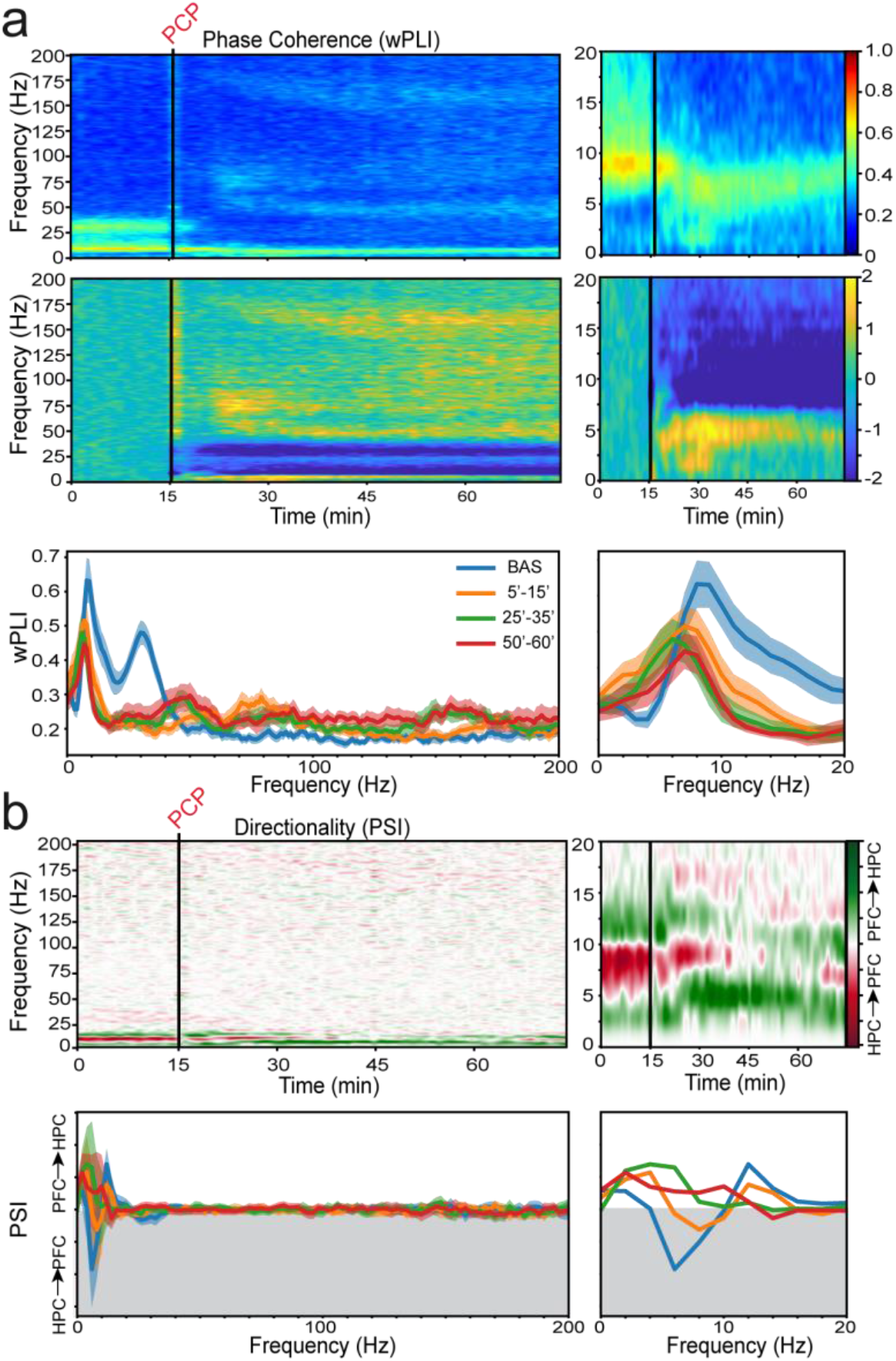
Acute administration of PCP disrupts prefrontal-hippocampal phase connectivity and directionality. **(a)** Time course of changes in weighted phase lag index (wPLI, phase coherence) and corresponding z-scores. Quantification of wPLI is shown below. **(b)** Time course of changes in phase slope index (PSI, circuit directionality). HPC-to-PFC directionality is represented in red and PFC-to-HPC directionality in green. A simplified representation of PSI is shown below where the shaded areas indicate HPC-to-PFC directionality (the line marks zero). 0-20 Hz magnifications are shown on the right. Average signals during baseline are shown in blue and signals during the 5-15, 25-35, and 50-60 min after PCP administration are represented in orange, green and red, respectively.

Major changes in circuit activity produced by PCP may be causally related to the mobility reduction it generated. To quantify this relationship, we analyzed periods of low mobility and compared them with periods of mobility at baseline (10 consecutive seconds in each state and mouse, n = 8 mice). Low mobility was determined by a defined threshold in the output of the accelerometer and baseline movement was defined as above that threshold. We found that PFC power was in fact sensitive to movement changes in baseline, specifically showing increased power during epochs of mobility compared to low mobility at theta, beta and gamma frequencies (F_1,7_ = 2.84, 3.62, 7.82, p = 0.025, 0.009, <0.0005). Moreover, as observed in the previous literature (Bohbot et al., 2017; Buzsáki, 2002), theta oscillations and theta-gamma coupling within the CA1 region were slightly increased during movement (F_1,6_ = 2.36, 2.18, p = 0.057, 0.072; Supplementary Fig. S2). In addition, during periods of low mobility there was a small increase in delta-gamma coupling in CA1 that was statistically insignificant (F_1,7_ = 2.08, p = 0.083). However, the circuit connectivity was overall not affected by the movement, only showing a significant increase in high gamma phase coherence (F_1,6_ = 2.84, p = 0.029). Following PCP administration, PFC theta, gamma power and high gamma connectivity increased despite the decreased mobility of the mice; therefore, these changes were likely due to the effects of PCP itself and not simply the increased immobility. However, PCP reductions of theta power and theta-gamma coupling in the HPC and beta power in the PFC were similar to decreases observed during epochs of immobility during baseline; therefore, these changes were likely influenced by the lower mobility of the mice after injection of PCP.

Collectively, these results suggest that PCP transforms the intrinsic HPC-to-PFC circuit synchronization at theta (8-12 Hz) into a PFC-to-HPC fast delta (4-8 Hz) mode. This pathological circuit reverberation at fast delta entrains alpha, high gamma and HFO in the PFC and high gamma and HFO in the HPC. This reorganization of the circuit dynamics interferes with local and circuit neural activity disrupting theta-gamma coordination. Over time, as synchronization at fast delta attenuates, the circuit becomes governed by high frequency oscillatory rhythms.

### Rescue of phencyclidine-induced prefrontal-hippocampal hypersynchronization and miscommunication by antipsychotic drugs

Antipsychotic medication including risperidone (RIS), clozapine (CLZ) and haloperidol (HAL) are prescribed to patients to stop or prevent psychotic symptoms. In addition, risperidone and clozapine may also ameliorate cognitive and negative symptoms. Therefore, it is important to understand how these two widely prescribed AAPDs affect prefrontal-hippocampal pathways, brain circuits relevant for cognition and depression (Godsil et al., 2013), and how they do so differently from the classical APD haloperidol. Fifteen minutes after PCP administration one of the three APDs was administered. We compared the neural signals recorded at 30 and 60 min after PCP injection between the group that only received PCP and each of the three PCP+APD groups. In these experiments, control injections with saline were performed before the administration of PCP.

We found that risperidone (2 mg/kg, I.P.; n = 6 mice) strongly inhibited all frequency bands in the PFC, reducing PCP-induced increases in power at fast delta, theta, alpha, high gamma and HFO (two-way ANOVA with time [baseline, 30 and 60 min after PCP] and treatment [PCP vs. PCP+RIS] as factors; interaction F_1,17_ = 4.81, 17.48, 6.01, 11, 11.82, p = 0.036, <0.0005, 0.02, 0.002, 0.001, respectively). In fact, theta, alpha, and high gamma power decreased below baseline levels ([baseline, saline, PCP+RIS] one-way ANOVA: F_1,5_ = 10.24, 12.39, 6.02, p = 0.001, <0.0005, 0.038; Fig. 3a,b). Risperidone did not reduce delta power but decreased its main frequency from fast to slow ranges (peak from 6.4 Hz to 3 Hz). In the HPC, risperidone recovered the power of PCP-induced HFO (F_1,16_ = 10.55, p <0.0005). Intriguingly, narrow bands in the high gamma and HFO ranges remained, steadily decreasing their frequency over time, suggesting that subpopulations of neurons still fired although their spike timing was highly restrained (Fig. 3a). In tandem with the fast delta and HFO power decreases, risperidone markedly reduced the PCP-induced fast delta-HFO PAC in the PFC (F_1,15_ = 5.91, p = 0.027), but the intrinsic theta-gamma coupling in the HPC was not restored (Fig. 3c). Similar effects were observed on the circuit PAC under the influence of risperidone. That is, anomalous inter-regional HPC_phase_-PFC_amp_ PAC subsided (F_1,15_ = 5.96, p = 0.028) but intrinsic PFC_phase_-HPC_amp_ theta-gamma coupling did not re-emerge (Fig. 3c). The inability of risperidone to reinstate inherent HPC theta oscillations and theta-gamma coupling could be influenced by the high immobility of mice (Acc: F_1,5_ = 36.52, p <0.0005, one-way ANOVA; Supplementary Fig. S2), which in fact remained more immobile under the effects of risperidone than PCP alone ([PCP+RIS vs. PCP alone] F_1,17_ = 145, p <0.0005, two-way ANOVA). Additionally, risperidone blocked the PCP-mediated increase in phase synchronization at high frequencies (F_1,16_ = 5.92, p = 0.028), however it was unable to rescue the alterations at lower frequencies (delta, theta, beta; Fig. 3d). Notably, it restored directionality of the circuit almost completely, redirecting the aberrant PFC-to-HPC fast delta flow of information towards the intrinsic HPC-to-PFC flow at theta (F_1,16_ = 5.93, p = 0.006; Fig. 3e). Overall, risperidone produced a very strong inhibition of neural activity and the animals’ behavior. We further investigated whether this was due to the dose used (2 mg/kg) or the nature of risperidone’s actions. A lower dose of 0.5 mg/kg produced very similar effects on mobility, coupling and connectivity, and only oscillatory power showed less suppression (Supplementary Fig. S3). We conclude that risperidone exerts strong inhibition of prefrontal-hippocampal circuits even at low doses.

**Figure 3.**
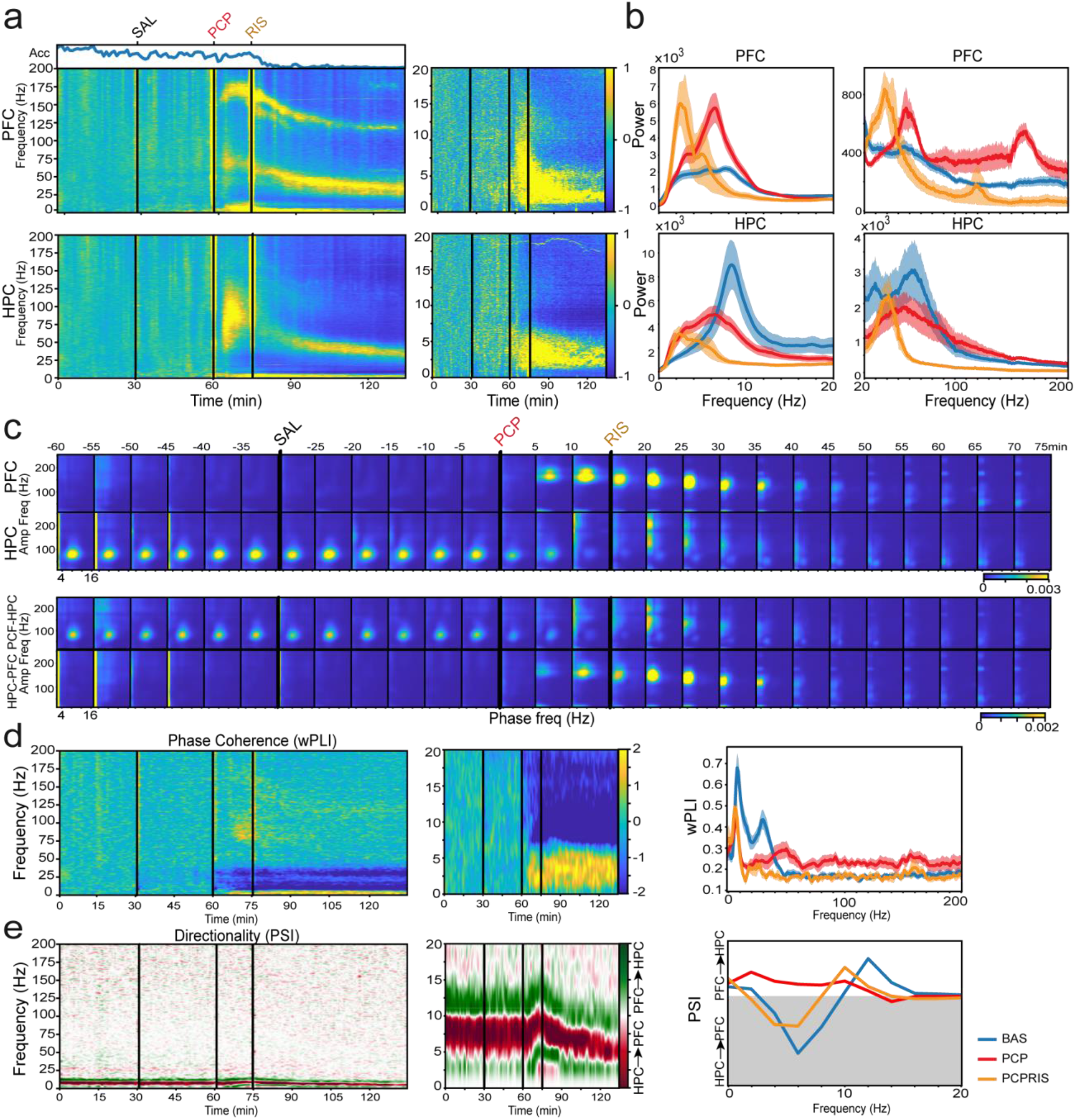
Risperidone (RIS) recovers prefrontal-hippocampal circuits by blocking PCP induced disruptions in power, cross-frequency coupling, connectivity, and directionality generated by PCP. **(a)** Normalized spectrograms (z-scores) of signals in the PFC (upper panels) and HPC (lower panels; n = 6 mice). The corresponding quantification of the animals’ mobility (Acc) is also shown. **(b)** The power spectra of PFC and HPC signals during 10 min of baseline are depicted in blue and signals from min 35 to 45 after RIS administration (50-60 min after PCP) are illustrated in orange. Note the pronounced shift of power by RIS from fast to slow delta in the PFC. An additional group with power spectra of signals recorded 50-60 minutes after the administration of PCP alone (the same as in figure 1d) are shown in red for comparison. **(c)** Comodulation maps quantifying cross-frequency coupling in consecutive non-overlapping 5-min epochs (as in Fig. 1e). The numbers on top of the panels indicate time after PCP administration. **(d)** Normalized (z-scores) time course of changes in wPLI (phase coherence) and corresponding quantification. **(e)** Normalized (z-scores) time course of changes in PSI (circuit directionality) and corresponding quantification. As above, the shaded area represents HPC-to-PFC directionality, the line denoting zero PSI. The group colors are the same as in b.

We next investigated whether clozapine (1 mg/kg, I.P.; n = 6 mice) blocked the actions of PCP on prefrontal-hippocampal neural dynamics, following the same experimental protocol as with risperidone (Fig. 4a). In general, clozapine reduced many PCP-induced power increases causing fewer inhibitory effects than risperidone. It decreased amplified theta and HFO power ([PCP+CLZ vs. PCP alone] power PFC theta and HFO: F_1,16_ = 3.78, 5.42, p = 0.033, 0.009; HPC HFO: F_1,16_ = 6.29, p = 0.005; two-way ANOVA) while aberrant alpha oscillations were partially rescued (F_1,16_ = 3.11, p = 0.057; Fig. 4a,b). Similar to risperidone, a narrow HFO band remained in the PFC. Also similar to risperidone, clozapine decreased the frequency of fast delta to slow delta (peak from 6.4 Hz to 3.14 Hz), blocked the *de novo* fast delta-HFO coupling (PFC, circuit: F_1,15_ = 3.77, 3.27, p = 0.054, 0.052), but was unable to reinstate baseline theta-gamma coupling in the CA1 and in the PFC-HPC circuit (Fig. 4c). The latter likely corresponds with the mild recovery in mobility after administration of clozapine (Acc: F_1,5_ = 6.82, p = 0.004). Functional connectivity alterations elicited by PCP were remarkably corrected by clozapine. First, PCP-strengthened phase coherence at low and high frequencies was normalized (fast delta, high gamma, HFO: F_1,16_ = 3.65, 5.143, 6.24; p = 0.059, 0.038, 0.026), however theta and beta coherence remained very low (Fig. 4d). Moreover, HPC-to-PFC flow of information at theta was fully rescued by clozapine (PSI: F_1,16_ = 6.180, p = 0.024), highly resembling baseline levels (p = 0.5; Fig. 4e).

**Figure 4.**
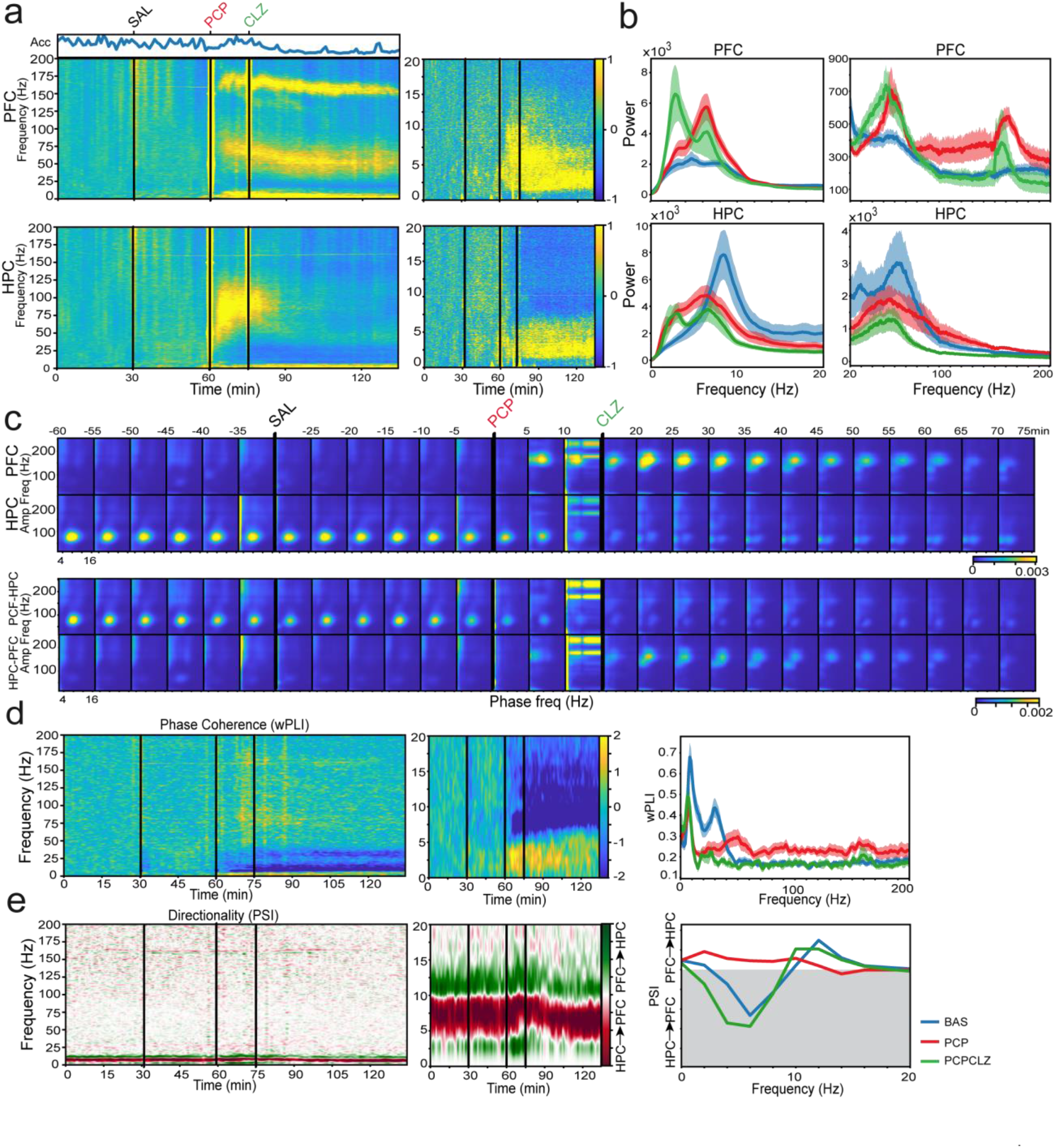
Clozapine (CLZ) blocks PCP-induced changes in power, cross-frequency coupling, circuit connectivity and directionality, generating less of an inhibitory effect compared to risperidone. **(a)** Normalized spectrograms (z-scores) of signals in the PFC (upper panels) and HPC (lower panels; n = 6 mice). The corresponding quantification of the animals’ mobility (Acc) is also shown. **(b)** The power spectra of PFC and HPC signals during 10 min of baseline are depicted in blue and signals from min 35 to 45 after CLZ administration (50-60 min after PCP) are illustrated in light green. As RIS, CLZ shifted the main power from fast to slow delta in the PFC. An additional group with power spectra of signals recorded 50-60 minutes after the administration of PCP alone (the same as in figure 1d) are shown in red for comparison. **(c)** Comodulation maps quantifying cross-frequency coupling in consecutive non-overlapping 5-min epochs (as in Fig. 1e). The numbers on top of the panels indicate time after PCP administration. **(d)** Normalized (z-scores) time course of changes in wPLI (phase coherence) and corresponding quantification. **(e)** Normalized (z-scores) time course of changes in PSI (circuit directionality) and corresponding quantification. Consistent with the figures above, the shaded area represents HPC-to-PFC directionality, the line denoting zero PSI. The group colors are the same as in **b**.

We finally examined the effects of the classical APD haloperidol (0.5 mg/kg, I.P.: n = 6 mice) on PCP-induced alterations (Fig. 5a). Haloperidol blocked all the aberrant bands elicited by PCP in the PFC except high gamma ([PCP+HAL vs. PCP alone] power fast delta, theta, alpha and HFOs: F_1,17_ = 6.74, 12.3, 5.48, 4.35, p = 0.014, 0.003, 0.032, 0.038; two-way ANOVA). In the HPC, it restored theta power (F_1,15_ = 6.1, p = 0.026), but was unable to rescue beta and HFO power changes. In addition, haloperidol reduced PFC and circuit fast delta-HFO coupling (F_1,14_ = 10.26, 6.78, p = 0.002, 0.008) and partly restored CA1 and circuit theta-gamma coupling (F_1,4_ = 2.79, 3.99, p = 0.086, 0.035, one-WAY ANOVA; not significantly different than PCP, two-WAY ANOVA; Fig. 5c). Hippocampal recovery of theta power and theta-gamma coupling was likely influenced by the fact that haloperidol recovered mobility more than risperidone and clozapine (Acc: F_1,5_ = 0.59, p = 0.63; one-WAY ANOVA). Interestingly, haloperidol produced novel cross-frequency coupling between slow delta (2-5 Hz) and HFO (100-140 Hz) in the HPC that could be also observed when it was injected alone (F_1,6_ = 10.2, p = 0.018; data not shown). Haloperidol also fully rescued phase coherence at low and high frequencies (fast delta and HFO: F_1,15_ = 5.98, 5.58, p = 0.027, 0.02) while failing to correct it at middle frequencies (alpha, beta and low gamma; Fig. 5d). Finally, haloperidol recovered HPC-to-PFC theta directionality (F_1,16_ = 6.25, p = 0.005; Fig. 5e).

**Figure 5.**
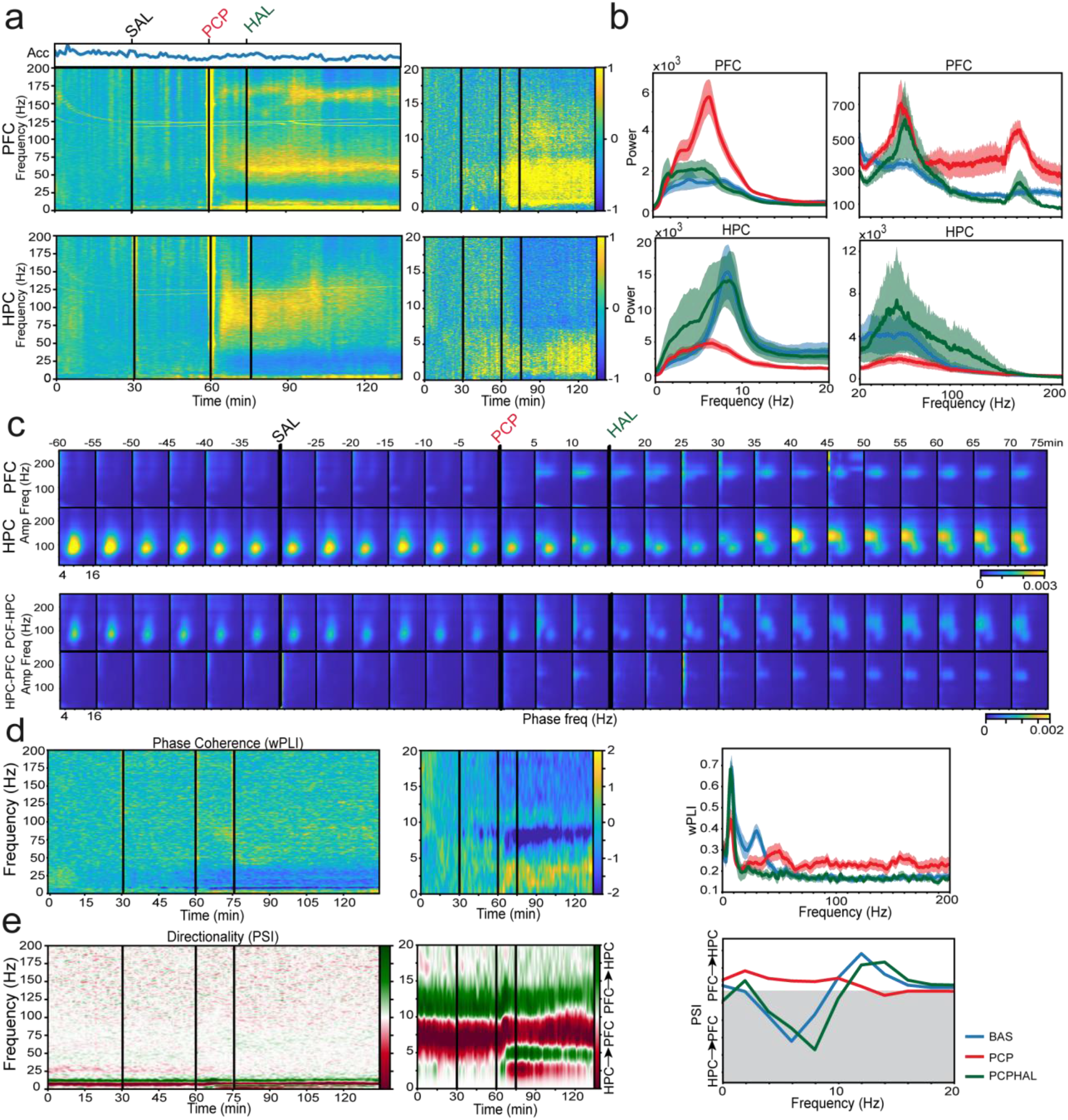
Haloperidol (HAL) rescues PCP-increased power, cross-frequency coupling and phase coherence and redirects abnormal circuit directionality. **(a)** Normalized spectrograms (z-scores) of signals in the PFC (upper panels) and HPC (lower panels; n = 6 mice). The corresponding quantification of the animals’ mobility (Acc) is also shown. **(b)** The power spectra of PFC and HPC signals during 10 min of baseline are depicted in blue and signals from min 35 to 45 after CLZ administration (50-60 min after PCP) are illustrated in dark green. An additional group with power spectra of signals recorded 50-60 minutes after the administration of PCP alone (the same as in figure 1d) are shown in red for comparison. **(c)** Comodulation maps quantifying cross-frequency coupling in consecutive non-overlapping 5-min epochs (as in Fig. 1e). The numbers on top of the panels indicate time after PCP administration. **(d)** Normalized (z-scores) time course of changes in wPLI (phase coherence) and corresponding quantification. **(e)** Normalized (z-scores) time course of changes in PSI (circuit directionality) and corresponding quantification. As above, the shaded area represents HPC-to-PFC directionality, the line denoting zero PSI. The group colors are the same as in b.

Together, the three APDs reversed many neurophysiological alterations produced by PCP, unraveling common cellular mechanisms of action. They were efficacious in correcting the aberrant power, cross-frequency coupling, phase coherence and directionality of signals generated by PCP. However, several differences also highlight that the cellular mechanisms varied between the three APDs. Overall, the AAPDs, risperidone and clozapine, were more inhibitory than haloperidol, both in general mobility of the animals and neural activity, which cannot be merely due to the higher doses used.

### Serotonergic receptors 5-HT_2A_ and 5-HT_1A_ play complementary roles in shaping prefrontal-hippocampal neural dynamics during phencyclidine-induced psychosis

We investigated the main serotonergic activities that could underlie the antipsychotic actions of risperidone and clozapine. Both compounds block 5-HT_2A_R directly and stimulate 5-HT_1A_R indirectly (Celada et al., 2013; Meltzer and Massey, 2011). Accordingly, we examined the abilities of a 5-HT_2A_R antagonist and a 5-HT_1A_R agonist to rescue PCP-induced prefrontal-hippocampal hypersynchronization. As mentioned above, we injected the drugs 15 minutes after PCP and compared the neural signals recorded at 30- and 60- minutes following administration of PCP between groups only given PCP and each of the PCP+5-HTR groups.

We first assessed the effects of the 5-HT_2A_R selective antagonist M100907 after PCP administration (1 mg/kg, I.P.; n = 6 mice). 5-HT_2A_R antagonism reduced PCP-generated bands in the PFC and the HPC ([PCP+M100907 vs. PCP alone] PFC fast delta, theta, alpha, high gamma, HFOs: F_1,17_ = 5.96, 9.44, 7.55, 4.43, 8.44, p = 0.026, 0.001, 0.014, 0.042, 0.001; HPC HFO: F_1,15_ = 8.46, p = 0.007, two-way ANOVA; Fig. 6a,b). Concurrent with the AAPDs, M100907 shifted the power of delta from the fast to the slow frequencies. Remarkably, narrow bands in the gamma and HFO ranges remained in both structures, following similar temporal patterns as risperidone. In addition, M100907 blocked increases in PFC high gamma and HFO power induced by the 5-HT_2A/2C_R agonist and hallucinogen DOI in the absence of PCP (1 mg/kg, IP; n = 7 mice; F_1,5_ = 16.55, 10.54, p = 0.001, < 0.0005; data not shown), which clearly indicates that 5-HT_2A_Rs are involved in shaping gamma and HFO rhythms in the PFC. 5-HT_2A_R antagonism also decreased PCP-induced fast delta-HFO coupling in the PFC and the circuit (F_1,14_ = 4.54, 5.52, p = 0.049, 0.01), while intrinsic theta-gamma coupling in CA1 and across regions shifted to delta-high gamma (2-5 to 50-80 Hz; F_1,14_ = 9.17, 10.74, p = 0.009, 0.006; Fig. 6c). This type of coupling is present in resting states during periods of little to no movement (Supplementary Fig. S2), therefore we hypothesize that this shift in coupling could be caused by a decrease in mobility after M100907 injection ([baseline, saline, PCP+M100907] Acc: F_1,5_ = 57.47, p <0.0005, one-way ANOVA; [PCP+M100907, PCP alone] F_1,17_ = 0.84, p = 0.41, two-way ANOVA). M100907 also partly normalized circuit phase hypersynchronization at high frequencies (wPLI gamma and HFO: F_1,15_ = 3.72, 6.08, p = 0.073, 0.006) but it did not affect the decreased synchronization at middle frequencies (Fig. 6d). To note, M100907 shifted fast delta to slow delta phase synchronization (F_1,15_ = 5.12, p = 0.039), closely following changes in delta power. Finally, HPC-to-PFC theta directionality was not recovered by M100907 (Fig. 6e). These results point to a major role of 5-HT_2A_R in AAPDs’ ability to reduce the PCP generated increases in power, cross-frequency coupling, and phase coherence, while not contributing to the normalization of the circuit’s directionality. 5-HT_2A_R antagonism may also be responsible for some immobility caused by AAPDs, as they show similar effects on theta power and theta-gamma coupling in the HPC.

**Figure 6.**
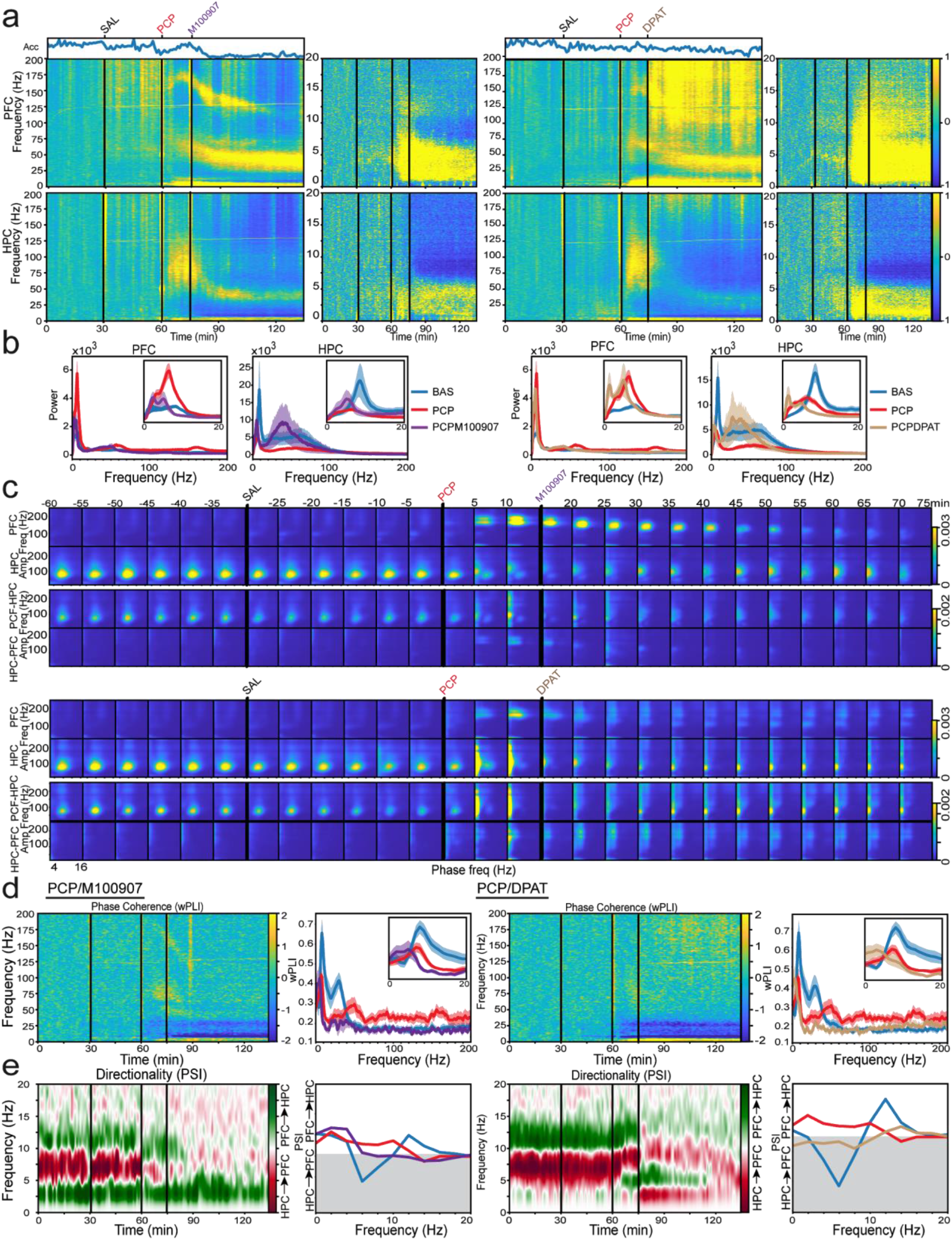
M100907 (5-HT_2A_R antagonist) and 8-OH-DPAT (DPAT, 5-HT_1A_R agonist) reverse different PCP-induced alterations of prefrontal-hippocampal neural dynamics. **(a)** The effects of M100907 are depicted on the left panels while the effects of DPAT are depicted on the right panels. Normalized spectrograms (z-scores) of signals in the PFC (upper panels) and HPC (lower panels; n = 6 mice each group). The corresponding quantification of the animals’ mobility (Acc) is also shown. **(b)** The power spectra of PFC and HPC signals during 10 min of baseline are represented in blue and signals from min 35 to 45 after M100907 or DPAT administration (50-60 min after PCP) are illustrated in purple and tan, respectively. Consistent with the figure above, an additional group with power spectra of signals recorded 50-60 minutes after the administration of PCP alone (the same as in figure 1d) are shown in red for comparison. **(c)** Comodulation maps quantifying cross-frequency coupling in consecutive non-overlapping 5-min epochs (as in Fig. 1e). The numbers on top of the panels indicate time after PCP administration. The effects of M100907 are depicted on the top panels while the effects of DPAT are depicted on the bottom panels. **(d)** Normalized (z-scores) time course of changes in wPLI (phase coherence) and corresponding quantification. **(e)** Normalized (z-scores) time course of changes in PSI (circuit directionality) and corresponding quantification. The shaded area represents HPC-to-PFC directionality, the line denoting zero PSI. The group colors are the same as in b.

We finally examined the contribution of 5-HT_1A_Rs to the modulation of prefrontal-hippocampal neural dynamics after injection of PCP. The selective 5-HT_1A_R agonist 8-OH-DPAT (DPAT, 1 mg/kg, I.P.; n = 6 mice) rescued power changes elicited by PCP in both brain regions, although it had less influence on prefrontal activities compared to M100907 (Fig. 6a,b). DPAT decreased power of fast delta, theta and high gamma oscillations in the PFC ([PCP+DPAT vs. PCP alone] F_1,17_ = 4.93, 9.09, 10, p = 0.04, 0.008, 0.003; two-way ANOVA), while HFOs were only slightly reduced compared mice only given PCP (F_1,17_ = 4.99, p = 0.014; [baseline, saline, PCP+DPAT] HFO: F_1,5_ = 6.28, p = 0.006; one-way ANOVA). In addition, there was only a partial transition from fast to slow delta power. In the HPC, PCP-altered beta and HFOs were fully corrected by DPAT (F_1,16_ = 6.08, 11.47, p = 0.015, 0.002; Fig. 6a,b). Notably, a prior study we conducted showed that DPAT given alone, in the absence of PCP, suppressed the power of high gamma and HFOs in both the PFC and HPC and such decreases were reversed by the selective 5-HT_1A_R antagonist WAY-100635 (PFC high gamma, HFO: F_1,6_ = 10, 9.12 p = 0.003, 0.001; HPC high gamma, HFO: F_1,6_ = 12.45, 21.66 p = 0.003, 0.001; one-way ANOVA; n = 7 mice)(Gener et al., 2019). This indicates that 5-HT_1A_Rs indeed play a role in shaping high gamma and HFO within the PFC and the HPC. Moreover, DPAT fully rescued increased fast delta-HFO coupling in the PFC and the entire circuit (F_1,16_ = 8.93, 3.37, p = 0.003, 0.049) while baseline CA1 and circuit theta-gamma coupling was not restored (Fig. 6c). This finding is quite notable because mice mobility was reduced less by DPAT than by M100907 (Acc: F_1,5_ = 4.38, p = 0.021, one-way ANOVA; [PCP+DPAT vs PCP+M100907] F_1,10_ = 5.66, p = 0.011, two-way ANOVA). In addition, 5-HT_1A_R agonism had limited effects on phase coherence and circuit directionality, corresponding to M100907’s actions. DPAT partially restored HFO phase connectivity (F_1,14_ = 3.14, p = 0.098), shifted phase synchronization from fast delta to slow delta (F_1,15_ = 4.99, p = 0.041) and did not recover HPC-to-PFC theta directionality (Fig. 6d,e). Thus, 5-HT_1A_R agonism could replicate some of the actions that AAPDs exerted on PCP-induced aberrant power and cross-frequency coupling, while its contribution to the normalization of the circuit’s directionality was probably minor.

## DISCUSSION

We found that the psychotomimetic effects of PCP were associated with hypersynchronization and disordered communication of prefrontal-hippocampal pathways. PCP induced major alterations in the medial PFC: it boosted oscillatory power at atypical frequencies within delta, alpha, high gamma and HFO ranges and generated fast delta-HFO coupling that suggested the presence of hypersynchronous cortical microcircuits. Cross-regional coupling and phase coherence at fast delta and HFOs were also enhanced, reflecting increased functional connectivity of the circuit. PCP also diverted intrinsic HPC-to-PFC theta signals into fast delta rhythms traveling in the opposite direction, from the PFC to the HPC. The time course of the changes suggests that prefrontal-hippocampal circuits were initially governed by fast delta rhythms following the administration of PCP. The circuit later transitioned into a state dominated by rhythms at high gamma and higher frequencies (<100 Hz) while the delta drive was attenuated. These events disrupted the normal function of the circuit, weakening intrinsic local and cross-regional theta-gamma coupling. The three APDs tested, risperidone, clozapine and haloperidol, reduced PCP-generated increase in power, cross-frequency coupling and phase coherence. Notably, they also reinstated theta directionality of the circuit, a key aspect considering that disrupted circuit communication is considered to be a hallmark of schizophrenia (Godsil et al., 2013; Hunt et al., 2017).

Our findings also support the proposed mechanism of a large-scale delta connectivity in schizophrenia, whereby an excessive thalamo-cortical delta drive disrupts the relationships between delta and higher frequencies in the thalamus and its downstream structures generating alterations in inter-regional connectivity (Hunt et al., 2017; Hunt and Kasicki, 2013). In fact, dissociative behaviors generated by NMDAR antagonists have been recently shown to correlate with slow oscillations within the delta range (1-3 Hz) that originate in the retrosplenial cortex (Vesuna et al., 2020). The consistent increase of delta power, coupling and connectivity induced by PCP occurred at standard “slow” delta frequencies in mice, between 2 to 5 cycles per second, but also at atypical fast delta frequencies, between 4 and 8 cycles per second, referred to as theta2 oscillations in rats (Ahnaou et al., 2017). All three APDs reduced the fast delta power generated by PCP, however the cellular mechanisms underlying this reduction were different for each APD. Both AAPDs, but not haloperidol, shifted the dominant frequency of delta oscillations from fast to slow ranges. The same effect was also produced by the two serotonergic compounds, which implies that changes in serotonergic transmission may have influenced the AAPDs’ shift in delta. Delta oscillations may be linked to lethargy and sedation observed in humans and animals treated with APDs, which would account for the significant decrease in the animals’ mobility observed after APD administration. Sedation has been attributed to APDs’ affinity for histamine H_1_ receptors (Miller, 2004). However, a recent study from our group also indicated that sedative effects may result from the cooperation between 5-HT_2A_/D2R antagonism and 5-HT_1A_ agonism (Gener et al., 2019).

Fast delta synchronization produced by PCP entrained aberrant network activities at high gamma and HFOs in the PFC and HPC, consequently boosting their power and circuit synchronization. Cortical gamma increases are observed during psychosis in healthy individuals and patients diagnosed with schizophrenia and are subsequently recovered by APDs (Jones et al., 2012; Shaw et al., 2015; Uhlhaas and Singer, 2010). In rodents, NMDAR antagonists such as PCP also increase gamma oscillations in cortical regions and this increase can also be rescued by APDs (Ahnaou et al., 2017; Caixeta et al., 2013; Hunt et al., 2017; Phillips et al., 2012). We found that risperidone, the 5-HT_2A_R antagonist, and the 5-HT_1A_R agonist clearly reversed the PCP-mediated increase of high gamma power in the PFC, while clozapine and haloperidol did not. This finding suggests 5-HT_2A_R antagonism and 5-HT_1A_R agonism play causal roles in risperidone’s ability to attenuate aberrant high gamma power in the PFC. Furthermore, our observations are consistent with findings from prior studies in rodents which have reported that risperidone blocks pathological high gamma oscillations more than clozapine (Ahnaou et al., 2017). To our knowledge, the presence of HFOs at the frequencies used here have been investigated in humans in the context of epilepsy but not psychosis, so an association between psychosis and HFOs still needs to be established. In rodents, the NMDAR antagonists PCP and ketamine produce rapid and substantial increases in the power of HFOs in several brain regions including cortical areas, the HPC, and the nucleus accumbens (Hunt et al., 2017, 2015). Accordingly, we also found that PCP produced significant increases in HFO power. HFO power was subsequently reduced by clozapine and risperidone in both the PFC and the HPC. However, the classical APD, haloperidol, was only able to reduce HFOs in the PFC and not the HPC. Additionally, M100907 and 8-OH-DPAT rescued PCP-induced HFOs in the HPC, which suggests that 5-HT_2A_R antagonism and 5-HT_1A_R agonism may mediate AAPDs’ unique ability to recover HFOs in the HPC. Finally, PCP also promoted prefrontal-hippocampal phase synchronization at high gamma and HFOs. This is consistent with the observed increases of phase coherence at gamma and HFOs across hippocampal layers after administration of ketamine in rodents (Caixeta et al., 2013). Notably, the phase synchronization was recovered by the three APDs and the two serotonergic drugs.

Previous studies suggest that abnormal activity of parvalbumin-expressing (PV+) interneurons may be responsible for the increase of gamma oscillations in schizophrenia. In support of this, postmortem studies of schizophrenia patients report reduced GABA synthesis in PV+ neurons (Gonzalez-Burgos et al., 2015). Moreover, mice lacking NMDARs on PV+ neurons have increased spontaneous gamma power in the hippocampus and cortex (Carlén et al., 2012; Korotkova et al., 2010). Therefore, in our study the blockade of NMDARs on PV+ neurons by PCP may have been critically involved in the amplification of high gamma oscillations. Importantly, different subpopulations of PV+ neurons in the PFC express 5-HT_1A_R and 5-HT_2A_R (Puig et al., 2010), rendering these neurons key elements in the gamma reduction mediated by AAPDs. In contrast, the neural mechanisms underlying PCP-mediated HFO are not as clear. A recent study has found a close correlation between the firing rates of PV+ neurons and HFOs in the medial PFC of healthy animals, although a causal relationship could not be established (Yao et al., 2020). Moreover, local blockade of NMDARs in the nucleus accumbens increases the power of HFOs (Hunt et al., 2010), although the neurons involved in generating these HFOs are unknown. Thus, the exact neural mechanisms underlying HFOs in the PFC and HPC that appear during psychotic states and their response to APDs will need further investigation.

In parallel with the circuit hypersynchronization, PCP persistently disrupted activity at theta, beta and low gamma which was not rescued by any of the APDs. PCP also disorganized the intrinsic hippocampal theta-gamma coupling, which was recovered by haloperidol, but not by the two AAPDs. As suggested by our control experiments in which we compared neural dynamics during epochs of movement and no movement, the immobility of mice caused by the AAPDs’ sedative effects may have contributed to such results. This is further supported by haloperidol’s greater ability to recover mobility compared to the AAPDs.

As mentioned above, our results suggest 5-HT_2A_R antagonism and 5-HT_1A_R agonism may have contributed to the actions of risperidone and clozapine. In particular, the way in which the 5-HT_2A_R antagonist, M100907, and risperidone blocked PCP’s actions appeared to be similar, which likely reflects risperidone’s strong affinity for 5-HT_2A_Rs. Both serotonergic compounds blocked the increase in power generated by PCP, although M100907 did so more effectively in cortical microcircuits whereas DPAT was more effective in hippocampal microcircuits. These results support the idea that 5-HT_2A_Rs and 5-HT_1A_Rs play complementary roles in shaping prefrontal-hippocampal neural dynamics (Puig et al., 2010). In contrast, the two serotonergic compounds were unable to recover disrupted directionality of the circuit. These results highlight that a combination of pharmacological activities may be required for antipsychotic action.

In conclusion, during psychotomimetic conditions induced via acute NMDAR blockade by PCP, prefrontal-hippocampal circuits appeared excessively hypersynchronized and exhibited disordered communication, which disrupted the intrinsic neural dynamics of the circuit. Immediately following administration of PCP, the circuits were governed by robust synchronization at delta that later transitioned into a state dominated by rhythms at higher frequencies while delta was attenuated. All APDs effectively corrected the PCP-induced increase in synchronization and disrupted circuit connectivity, which may point to a common neural mechanism in how APDs treat the core symptoms of psychosis. However, differential effects were identified between the AAPDs and the classical APD, haloperidol, likely reflecting their distinct affinity for serotonin and dopamine receptors. We investigated the actions of APDs under the psychotic-like states generated by acute PCP. Understanding how APDs affect prefrontal-hippocampal neural dynamics in animal models of the cognitive and negative symptoms in schizophrenia (e.g., via subchronic PCP administration) is imperative to make progress in the development of novel therapeutic interventions for neuropsychiatric diseases.

## AUTHOR CONTRIBUTIONS

CDS, TG and MVP designed research; CDS, TG, MT and ABF performed research; CDS and PN analyzed data; all the authors wrote the paper.

## MATERIALS AND METHODS

### Animals

Adult male C57BL/6 mice (n = 30) were obtained from the local colony at the Barcelona Biomedical Research Park (PRBB) animal facility. They were 2 to 3 months old at the start of the experiments and weighed between 20 and 30 g. All procedures were conducted in compliance with EU directive 2010/63/EU and Spanish guidelines (Laws 32/2007, 6/2013 and Real Decreto 53/2013) and were authorized by the PRBB Animal Research Ethics Committee and the local government (Generalitat de Catalunya). Mice were maintained under controlled environmental conditions throughout the study: 22 ± 2 °C ambient temperature, the relative humidity at 60%, 12:12 light–dark cycle (lights off from 07:30 p.m. to 7:30 a.m.), food (standard pellet diet) and water available *ad libitum*.

### Surgeries

We used similar procedures to those in Gener et al., 2019. Briefly, mice were induced with ketamine/xylazine and anesthesia was maintained with isoflurane at 0.5-4%. Several micro-screws were placed into the skull to stabilize the implant, and the one on top of the cerebellum was used as a general ground. Three tungsten electrodes (25 μm wide, 100 to 400 kOhm; Advent, UK), one stereotrode and one single electrode, were unilaterally implanted in the prelimbic PFC (AP: 1.5, 2.1 mm; ML: ± 0.6, 0.25 mm; DV: −1.7 mm from bregma) and two more were implanted in the CA1 area of the HPC (AP: −1.8 mm; ML: −1.3 mm; DV: −1.15 mm). After surgery animals were allowed at least one week to recover during which they familiarized with the implant connected to the recording cable. After the experiments ended, the electrode placements were confirmed histologically by staining the brain slices (30-μm-thick coronal slices using a cryostat) using Cresyl violet. Images were obtained and analyzed using light microscopy (Olympus BX61) with a 4x magnification. Electrodes with tips outside the targeted areas were discarded from data analyses.

### Neurophysiological recordings and data analyses

Electrophysiological recordings were conducted between 9:00 a.m. and 5:00 p.m., during the light cycle of housing, in freely-moving mice exploring their home cages (369 × 165 × 132 mm). Animals did not have access to food or water during the recording sessions but had food and water available just before and after each experiment. All the recordings were carried out with the multi-channel Open Ephys system at 0.1-6000 Hz and a sampling rate of 30 kHz with Intan RHD2132 amplifiers equipped with an accelerometer. The home cage was moved from the housing room to the experimental room within the animal facility where recordings were implemented. Two to three animals were recorded simultaneously in separate cages and electrophysiological setups in the same room separated by partitions.

Recorded signals from each electrode were detrended, notch-filtered to remove power line artifacts (50, 100, 150 and 200 Hz) and decimated to 1kHz offline to obtain local field potentials (LFPs). Noisy electrodes detected by visual inspection from individual channel spectrograms were not used. Power spectral density results were calculated using the multi-taper method from the spectral_connectivity package in Python (time-half-bandwidth product = 5, 9 tapers, 60s sliding time window without overlap; https://github.com/Eden-Kramer-Lab/spectral_connectivity). Spectrograms were constructed using consecutive Fourier transforms (scipy.signal.spectrogram function, 60s time window, no overlap, no detrend). A 1/f normalization was applied to power spectral density results, and power spectrograms were scaled to decibels for visualization purposes. The frequency bands considered for the band-specific analyses included: delta (2-5 Hz), fast delta (4-8 Hz), theta (8-12 Hz), alpha (10-14 Hz), beta (18-25 Hz), low gamma (30-48 Hz), high gamma (52-100 Hz), PCP-induced narrow high gamma (52-75 Hz), HFO (100-200 Hz) and PCP-induced narrow HFO (130-180 Hz). Phase-amplitude coupling (PAC) was measured with a Python implementation of the method described in Tort et al. (phase frequencies = [0, 15] with 1 Hz step and 4 Hz bandwidth, amplitude frequencies = [10, 250] with 5 Hz step and 10 Hz bandwidth; (Tort et al., 2008). The length of the sliding window was 300s for the overview plots and 60s for the quantifications, without overlap. PAC quantification results were obtained by averaging the values of selected areas of interest in the comodulograms. Prefrontal-hippocampal functional connectivity was estimated via the weighted phase-lag index (wPLI, Butterworth filter of order 3), a measure of phase synchronization between areas aimed at removing the contribution of common source zero-lag effects that allowed us to estimate the synchronization between the PFC and the HPC mitigating source signals affecting multiple regions simultaneously (Gener et al., 2019; Hardmeier et al., 2014; Stam et al., 2007; Vinck et al., 2011). PLI spectras were built applying the previous function multiple times with a 1 Hz sliding frequency window (using Butterworth bandpass filters of order 3), and PLI spectrograms were generated applying the PLI spectra function over a 60s sliding window (without overlap). In addition, we calculated the flow of information between areas with the phase slope index (PSI) with a Python translation of MATLAB’s data2psi.m (epleng = 60s, segleng = 1s) from (Nolte et al., 2008). PSI spectrograms and spectra were constructed with the same strategy as PLI plots but using a 2 Hz sliding frequency window. All the results (LFP power, local and circuit PAC and PFC-HPC wPLI) are provided as z-scores with respect to baseline statistics (i.e., data is demeaned by the baseline mean and then normalized by the baseline standard deviation).

We used the accelerometer’s signals to evaluate the effects of the drugs on general mobility of mice. We found that the variance of the acceleration module (Acc), which quantifies the variation of movement across the three spatial dimensions, was largest during exploration and decreased as the animals were in quiet alertness or sedation. More specifically, we calculated the instantaneous module of raw x, y and z signals from which we measured the variance of 1 minute bins (Gener et al., 2019). The Acc results were presented as a ratio to the highest value in the baseline condition of the respective experiment. Ten second epochs during quiet wakefulness and movement were identified applying a threshold to the accelerometer data as in (Alemany-González et al., 2020). Linear interpolation was applied to remove injection artifacts in Acc plots for visualization purposes (a single point was affected in each case).

### Pharmacology

Phencyclidine (noncompetitive NMDAR antagonist) was obtained from LGC STANDARDS S.L.U. Risperidone (RIS, atypical APD), clozapine (CLZ, atypical APD), haloperidol (HAL, D_2/3_ antagonist and typical APD), M100907 (5-HT_2A_ antagonist), 1-(2,5-dimethoxy-4-iodophenyl)-2 amino propane (DOI, partial 5-HT_2A/2C_R agonist), and 8-hydroxy-2-(di-n-propylamino)tetraline (8-OH-DPAT, 5-HT_1A_ agonist) were obtained from Sigma/Aldrich. WAY-100635 (WAY, 5-HT_1A_R antagonist) was obtained from Biogen Científica (Madrid, Spain). All drugs were dissolved in saline solution, except RIS that was dissolved in 1% DMSO solution. Doses were: PCP 10 mg/kg; RIS 2 and 0.5 mg/kg; CLZ 1 mg/kg; HAL 0.5 mg/kg; M100907 1 mg/kg; 8-OH-DPAT 1 mg/kg; DOI 1 mg/kg; WAY 0.5 mg/kg (Gener et al., 2019). All drugs were administered intraperitoneally (I.P.), with the exception of PCP, which was administered subcutaneously (S.C.). Each mouse was used to test multiple drugs and at least one week of washout was left between experiments.

### Experimental design and statistical analyses

The initial experiments consisted of 15 minute baseline periods and 1 hour periods after administration of PCP. In experiments that aimed to reverse the effects of PCP, additional 30 min control periods with saline were introduced after baseline, and then risperidone, clozapine, haloperidol, M100907 and 8-OH-DPAT were administered 15 minutes after PCP. One-way repeated measures ANOVA was used to test for statistical significance with time as within subject factor (baseline, saline, PCP and drug) for Acc, power, PAC, wPLI and PSI with IBM SPSS Statistics software (PASW Statistics 18). Additionally, two-way repeated measures ANOVA was used to test for statistical significance with treatment as between factor (PCP alone vs. saline controls and PCP alone vs. PCP+drug) and time as within subject factor (baseline, saline, PCP and drug). Drug effects were quantified in 10 min epochs (last 10 min of baseline and saline, minutes 25-35 and 50-60 after PCP injections). Raw data were used in all statistical analyses. We used the Bonferroni method to adjust for multiple comparisons.

