## Supplemental Information for "Phencyclidine-induced psychosis causes hypersynchronization and disruption of connectivity within prefrontal-hippocampal circuits that is rescued by antipsychotic drugs"

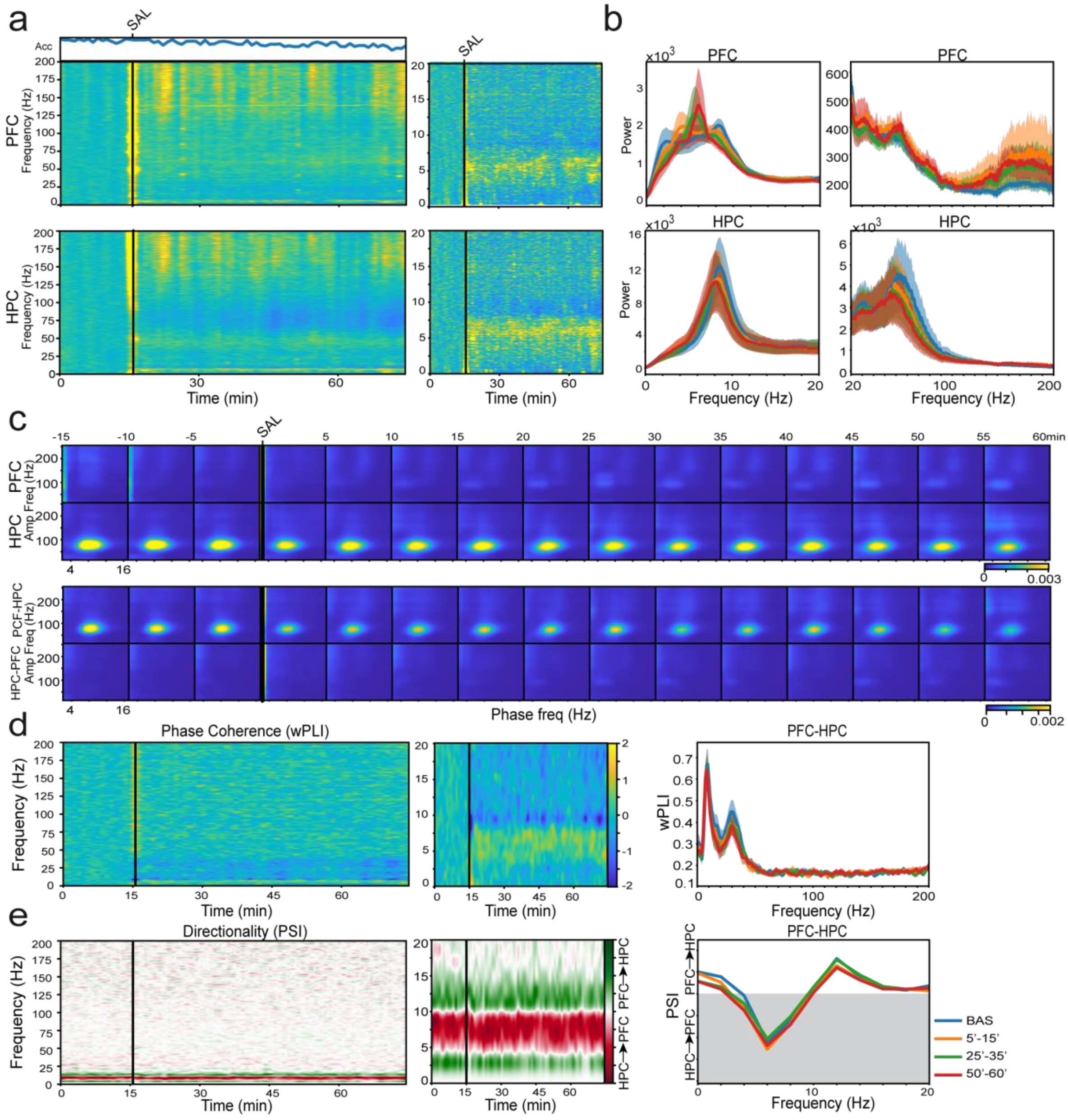

**Supplementary Figure S1.** Saline has minor effects on prefrontal-hippocampal neural dynamics. **(a)** Normalized spectrograms (z-scores) of signals in the PFC (upper panels) and HPC (lower panels; n = 10 mice). The corresponding quantification of the animals' mobility (Acc) is also shown. **(b)** Power spectra of PFC and HPC signals. The plots have been divided into 0-20 Hz and 20-200 Hz to facilitate visualization. **(c)** Comodulation maps quantifying cross-frequency coupling in consecutive non-overlapping 5-min epochs. **(d)** Normalized (z-scores) time course of changes in wPLI (phase coherence) and corresponding quantification. **(e)** Normalized (z-scores) time course of changes in PSI (circuit directionality) and corresponding quantification. The shaded area represents HPC-to-PFC directionality and the line indicates zero PSI. Group colors: baseline (blue), 5 to 15 min (orange), 25 to 35 min (green) and 50 to 60 min (red) after the administration of saline.

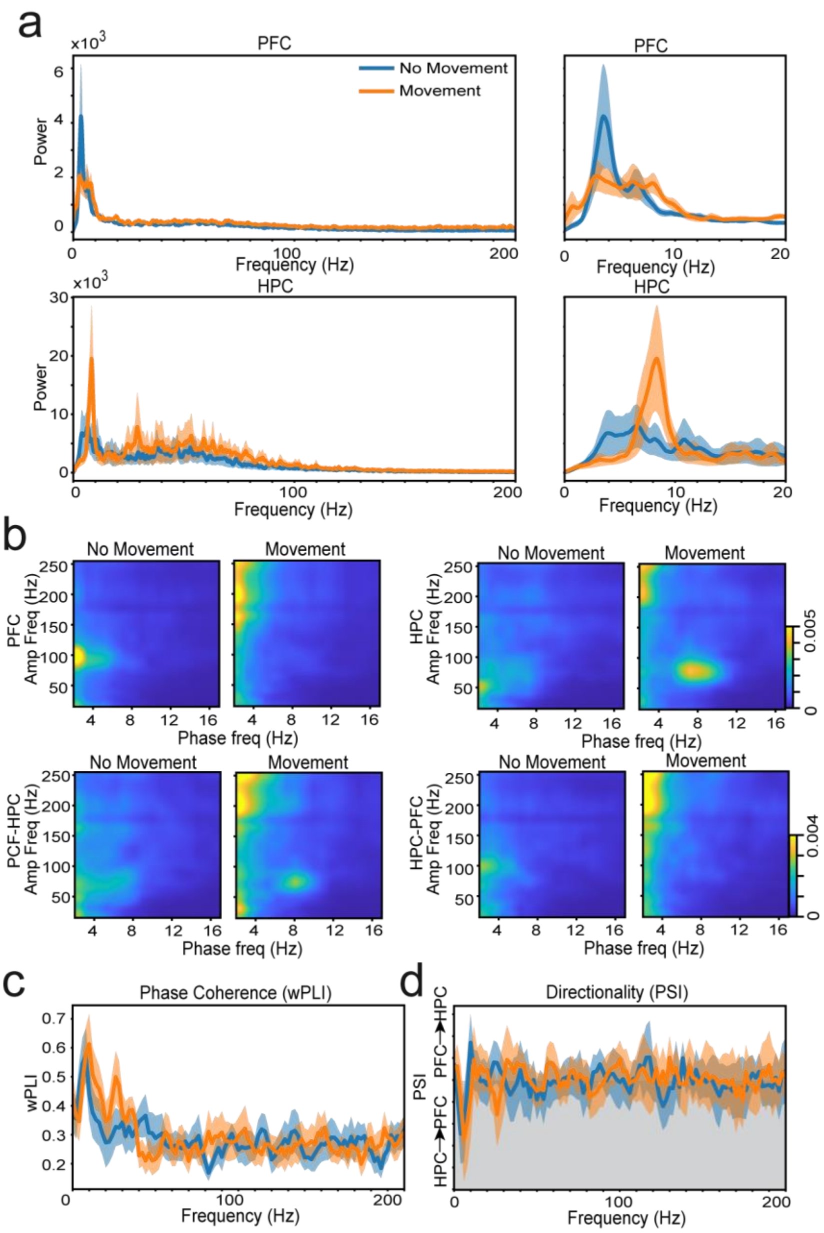


**Supplementary Figure S2.** Little influence of mice mobility in prefrontal-hippocampal neural dynamics. We analyzed periods of low mobility and compared them to periods of normal mobility during baseline (n = 8 mice, 10 consecutive second epochs). Low mobility was determined by a defined threshold in the output of the accelerometer signals and normal movement was defined as above that threshold. **(a)** Power spectra of PFC and HPC signals while animals rest or move. The plots have been divided into 0-20 Hz and 20-200 Hz to facilitate visualization. **(b)** Averaged comodulation maps of local and cross-regional modulation index in the two groups. **(c)** Average wPLI in the two groups. **(d)** Average PSI in the two groups. The shaded area represents HPC-to-PFC directionality and the line indicates zero PSI.

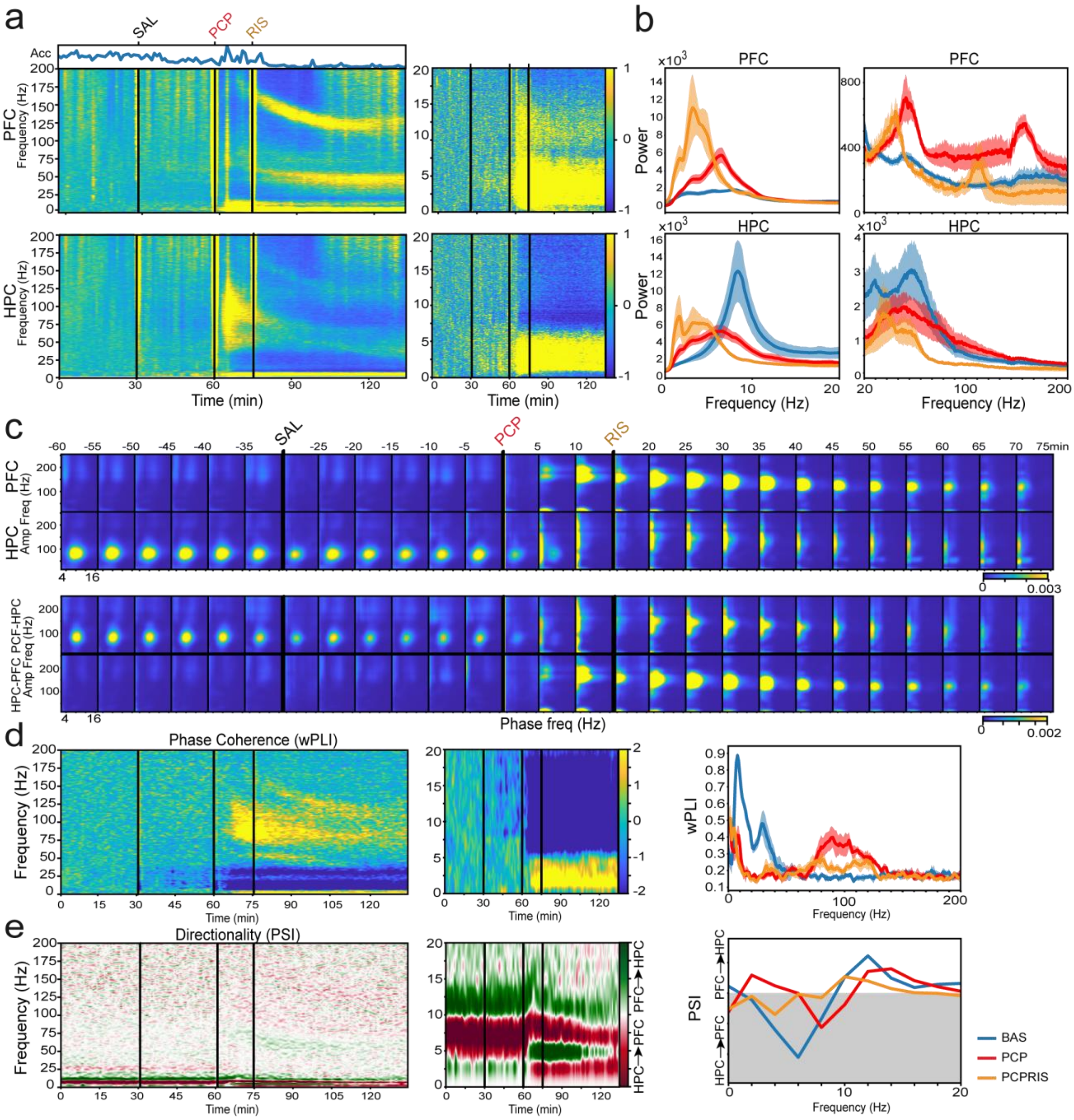
a
SAL
PCP
RIS
Acc
PFC Frequency (Hz)
HPC Frequency (Hz)
Time (min)
b
PFC
HPC
Power
Frequency (Hz)
c
PFC
HPC
Amp Freq (Hz)
HPC-PFC PCF-HPC
Phase freq (Hz)
75min
d
Phase Coherence (wPLI)
Frequency (Hz)
wPLI
e
Directionality (PSI)
PSI
PFC→HPC
HPC→PFC
BAS
PCP
PCPRIS

**Supplementary Figure S3.** A lower dose of risperidone (RIS; 0.5 mg/kg) causes inhibition of prefrontal-hippocampal circuits and blocks the effects of PCP. **(a)** Normalized spectrograms (z-scores) of signals in the PFC (upper panels) and HPC (lower panels; n = 3 mice). The corresponding quantification of the animals' mobility (Acc) is also shown. **(b)** The power spectra of PFC and HPC signals during 10 min of baseline are depicted in blue and signals from min 35 to 45 after RIS administration (50-60 min after PCP) are illustrated in orange. Note the impressive shift of power by RIS from fast to slow delta in the PFC. An additional group with power spectra of signals recorded 50-60 minutes after the administration of PCP alone (the same as in figure 1d) are shown in red for comparison. **(c)** Comodulation maps quantifying cross-frequency coupling in consecutive non-overlapping 5-min epochs. The numbers on top of the panels indicate time after PCP administration. **(d)** Normalized (z-scores) time course of changes in wPLI (phase coherence) and corresponding quantification. **(e)** Normalized (z-scores) time course of changes in PSI (circuit directionality) and corresponding quantification. As above, the shaded area represents HPC-to-PFC directionality, the line denoting zero PSI. The group colors are the same as in b.
